# Derivation and Validation of a Record Linkage Algorithm between EMS and the Emergency Department

**DOI:** 10.1101/124313

**Authors:** Colby Redfield, Abdulhakim Tlimat, Yoni Halpern, David Schoenfeld, Edward Ullman, David A Sontag, Larry A Nathanson, Steven Horng

**Affiliations:** Department of Emergency Medicine, Beth Israel Deaconess Medical Center, Boston, MA; Department of Computer Science, New York University, New York, New York

## Abstract

**Background:** Linking EMS electronic patient care reports (ePCRs) to ED records can provide clinicians access to vital information that can alter management. It can also create rich databases for research and quality improvement. Unfortunately, previous attempts at ePCR - ED record linkage have had limited success.

**Objective:** To derive and validate an automated record linkage algorithm between EMS ePCR’s and ED records using supervised machine learning.

**Methods:** All consecutive ePCR’s from a single EMS provider between June 2013 and June 2015 were included. A primary reviewer matched ePCR’s to a list of ED patients to create a gold standard. Age, gender, last name, first name, social security number (SSN), and date of birth (DOB) were extracted. Data was randomly split into 80%/20% training and test data sets. We derived missing indicators, identical indicators, edit distances, and percent differences. A multivariate logistic regression model was trained using 5k fold cross-validation, using label k-fold, L2 regularization, and class re-weighting.

**Results:** A total of 14,032 ePCRs were included in the study. Inter-rater reliability between the primary and secondary reviewer had a Kappa of 0.9. The algorithm had a sensitivity of 99.4%, a PPV of 99.9% and AUC of 0.99 in both the training and test sets. DOB match had the highest odd ratio of 16.9, followed by last name match (10.6). SSN match had an odds ratio of 3.8.

**Conclusions:** We were able to successfully derive and validate a probabilistic record linkage algorithm from a single EMS ePCR provider to our hospital EMR.

## 1. Introduction

### Background

Electronic patient care reports (ePCRs) by emergency medical services (EMS) providers yield critical information that improves patient care in the emergency department and throughout the entire hospital stay. Through these ePCRs, emergency medicine staff gain insight into the circumstances leading up to the emergency including information from bystanders on scene, the clinical trajectory of the patient, nature of prehospital treatments administered and other information which would otherwise be lost.^1^ The utility of prehospital documentation extends beyond emergency physicians to the interdisciplinary care team. Social workers, case managers, physical therapists and nurses can develop better care plans if they have information about the patient’s home situation, such as of the number of stairs they must climb, the safety and cleanliness of the surroundings and contributing factors such as substance abuse and domestic violence. The inpatient team can refer back to prehospital documentation to better correlate new facts that come to light and the patient’s response to treatment (or lack thereof).

### Importance

The ongoing transition from traditional paper records to ePCRs has been a critical transformation to allow inclusion of prehospital information into the hospital’s electronic medical record (EMR). However this integration has been hampered due to the lack of a reliable method to automatically link a patient’s data in the EMS computer to the correct patient record in the EMR. This type of record linkage, matching the same patient’s records across different data sources, is very challenging in the US healthcare system due to the lack of a national patient identifier. Combining data from different data sources increases the breadth and depth of information that can be analyzed.^2^ Record linkage has been attempted in prior studies for EMS records with limited success.^3^ Having accurate record linkage can serve many important functions in the emergency department for both prospective clinical use and retrospective data analysis. In the real-time clinical setting, linkage of the ePCR to the ED EMR provides the EM clinician access to vital information that can alter management in the ED. Studies have shown that physicians find PCRs to be important for Emergency Department care and medical decision making, and the lack of scene data is associated with increased risk of mortality in trauma patients.^4^ Retrospectively, the ability to match records accurately can improve data sets for research by augmenting existing clinical registries such as The National Trauma Data Bank and Cardiac Arrest Registry to Enhance Survival.^5^

In the past, the inability to match prehospital data with hospital outcomes has limited the utility of such data sets and potentially hindered advancements in trauma care, cardiac arrest, stroke, and prehospital research in general. Besides the utility in research, record linkage can promote further quality assurance and education to both EMS providers and ED providers. Previously, it was difficult for EMS providers to receive contemporaneous feedback about their individual patient outcomes. Having discharge diagnosis and information linked to an ePCR would allow EMS administration and individual providers to continuously improve their practice, target educational interventions, and automatically monitor key performance metrics.

### Goals of this Investigation

The goal of our study is to derive and validate a record linkage algorithm to accurately link EMS ePCRs to hospital EMR systems.

## 2. Methods

### Study Design and Setting

We conducted a retrospective derivation and internal validation study of a record linkage algorithm between EMS ePCRs and ED EMR’s using state-of-the-art machine learning techniques. The study was submitted to our institutional review board and a determination was made that no further review was required. The study was performed at a 55,000 visits/year Level I trauma center and tertiary academic teaching hospital.

### Selection of Participants

All consecutive adult ED patients who arrived in the ED between June 2013 and June 2015 from a single EMS agency were included. No patients were excluded. The hospital uses a locally developed ED information system (EDIS) known as the “ED Dashboard” which serves as the EMR for ED patients at this institution.

### Data Collection and Processing

Prehospital care occurs in a high pressure environment where providers must record information with limited time, incomplete data, and competing priorities. While errors may be present, we leverage modern machine learning techniques to maximize the value of what was captured, without requiring further manual review.

We electronically extracted 6 commonly found data elements from both the ePCR and EMR: age, gender, last name, first name, social security number (SSN), and date of birth (DOB). The features used in the dataset are listed in [Table 1].

**Table 01:** Model Features

| Feature | Description |
| --- | --- |
| Identical data features |  |
| *dob_match* | date of birth is identical in both records |
| *dob_levenshtien* | Levenshtein edit distance between date of birth in each record |
| *gender_match* | gender is identical in both records |
| *lastname_match* | last name is identical in both records* |
| *firstname_match* | first name is identical in both records* |
| *ssn_match* | SSN is identical in both records |
| Data comparison features |  |
| *age_diff* | difference in age between two records |
| *SSN_levenshtein* | Levenshtein edit distance between social security numbers in each record |
| *last_name_jw* | Jaro-Winkler edit distance between the two last names* |
| *first_name_jw* | Jaro-Winkler edit distance between the two first names* |
| Missing Data Indicators |  |
| *missing_emr_age* | missing variable indicator if age missing from ED data |
| *missing_emr_dob* | missing variable indicator if date of birth missing from ED data |
| *missing_emr_firstname* | missing variable indicator if first name missing from ED data |
| *missing_emr_lastname* | missing variable indicator if last name missing from ED data |
| *missing_emr_gender* | missing variable indicator if gender missing from ED data |
| *missing_emr_ssn* | missing variable indicator if social security number missing from ED data |
| *missing_ems_gender* | missing variable indicator if gender missing from ePCR |
| *missing_ems_firstname* | missing variable indicator if first name is missing from ePCR |
| *missing_ems_age* | missing variable indicator if age missing from ePCR |
| *missing_ems_dob* | missing variable indicator if date of birth is missing from ePCR |
| *missing_ems_ssn* | missing variable indicator if social security number missing from ePCR |
| *missing_ems_lastname* | missing variable indicator if last name missing from ePCR records |

\* First name – last name transposition are allowed without penalty.

Features were constructed to allow for common transposition errors. For example, the “*firstname_match*” and “*lastname_match*” features permit the first and last names to be swapped without penalty.

Typographic errors between the ePCR and EMR social security numbers were accounted for using the Levenshtein edit distance metric, which was then scaled to lie between 0 and 1.^6^ In the following example, the Levenshtein edit distance would be 2 because it would take 2 edits to transform the string “CAR” to “ARK”

1. <u>C</u>AR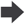 AR [1 deletion]
2. AR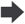 AR<u>K</u> [1 addition ]

Typographical errors in first and last names were accounted for using the Jaro-Winkler (J-W) distance^2^ metric, which is a measure that lies between 0 and 1, where 1 represents a perfect match. For example, we calculate the feature “firstname_jw” by finding the J-W distance between the EMR first name and the ePCR first name. We also calculate the J-W distance between the EMR first name and the ePCR last name in case the first and last names were accidentally swapped. We use the best (larger) of the two J-W distances.

The *“age_diff”* was calculated as a percentage error and then divided by 100 to scale the value to lie between 0 and 1, using the formula:

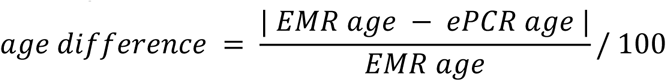

A missing indicator feature variable was created for age, gender, last name, first name, SSN and DOB for each of the EMR and ePCR datasets to denote missing data.

The data set was comprised of every pair of ePCR and EMR records that was registered within +/- 2 hours of the EMS arrival time indicated in the ePCR that is described in detail in [Figure 01].

**Figure 01:**
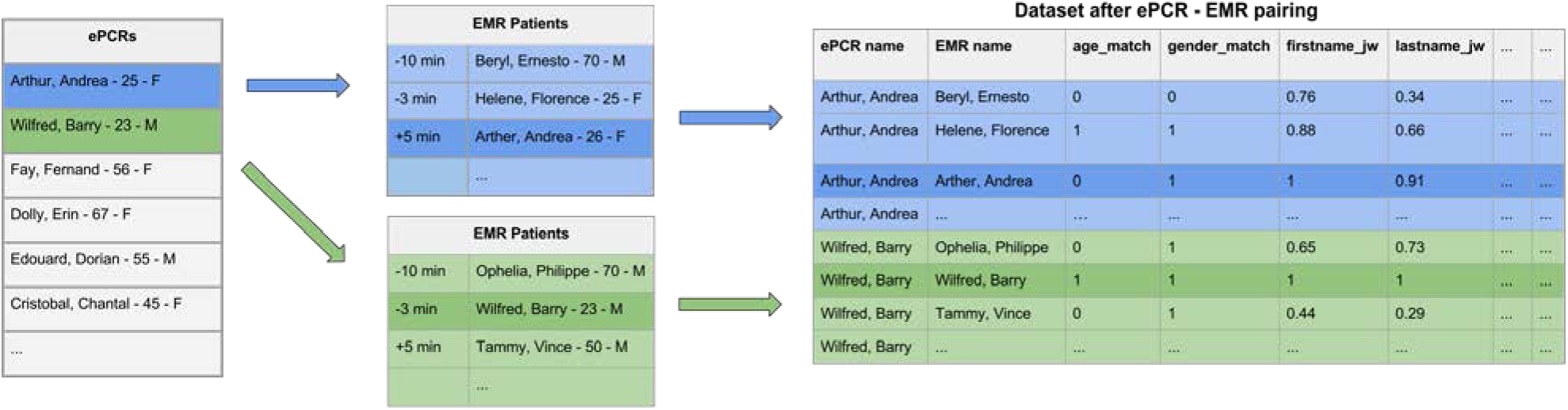
Dataset Generation

ePCR-EMR pairs were randomly allocated to a train (80%) or test (20%) data set, where the unit of randomization was at the level of the ePCR records to ensure that all sets of ePCR-EMR pairs were randomized to the same data set.

### Outcome Measure

A human reviewer manually matched records to determine a gold standard match between ePCRs and ED visits. For each ePCR, the primary reviewer was presented a time ordered list of ED visits that began within a 2 hour window around the reported time of EMS arrival. The reviewer matched each ePCR to an ED visit patient record to create a gold standard linkage. This linkage was based on the demographic fields from each source as discussed above (name, DOB, age, gender, and SSN) as well as additional data elements (arrival time, chief complaint, patient home address, the EMS historical narrative and the ED nursing triage summary). A positive example was defined if the reviewer linked an ePCR to that EMR record. A negative example was defined if the reviewer did not link the ePCR. Since there can only be one positive example per ePCR and many negative examples, there exists a class imbalance between positive and negative classes that is described below in the Model Derivation section.

In order to ensure that our labelling method was accurate, a second reviewer randomly oversampled the primary reviewer’s linkages (n=1400; 10%) and calculated a Cohen’s Kappa.

### Primary Data Analysis

Statistical analysis was performed using the python scikit-learn package.^7^ Means with 95% confidence intervals were reported for normally distributed variables and medians with interquartile ranges were reported for non-normal variables.

### Model Derivation and Analysis

We used the python scikit-learn package^7^ to train a multivariate logistic regression model using 5-K fold cross-validation on the train data set. We used the “label k-fold” feature of sci-kit learn’s cross-validation package to ensure that all ePCR pairs were assigned to the same data set. Given the large number of features, we used L2 regularization to prevent overfitting to fully utilize all the features, rather than an automated feature selection method. The optimal regularization parameter C was chosen using cross-validation. We also assigned sample weights to negative samples to account for the large class imbalance between positive and negative samples using the formula:

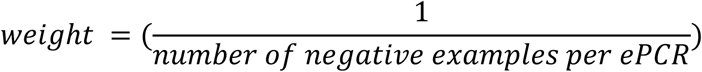

We report the sensitivity, positive predictive value, area under the receiver operating characteristic curve (AUC) on both the 80% training data set, and the 20% held out test data set.

## 3. Results

### Characteristics of Study Subjects

A total of 14,032 patients were enrolled during the study period. The mean age was 53.0, and 47.9% were males. For Urgency to Scene, 14% patients had priority 1 which represents a time sensitive or life threatening event, 13.4% patients had priority 2 which represents a non-life threatening event, 7.8% had priority 3 which represents non-acute injury, 0.2% patients had priority 4 which represents hold until verified need by another responding agency such as police or fire and 64.4% had an undocumented priority. For Urgency from Scene, 5.7% patients had priority 1, 26.8% patients had priority 2, 25.8% had priority 3 and 41.7% had an undocumented priority. The EMS skillset was ALS in 10.9% patients and BLS in 88.7%. 84.7% of the patient population spoke English, with a total of 39 languages. More detailed patient demographics are shown in [Table 2]

**Table 02:** Patient Demographics

|  |  |  |
| --- | --- | --- |
| Age - years (95% CI) | 53.0 (52.6 - 53.3) |  |
| Male gender - n (%) | 6724 (47.9%) |  |
| Urgency To Scene |  |  |
|  | Priority 1 (time sensitive or life threatening event) | 1976 (14%) |
|  | Priority 2 (non-life threatening event) | 1884 (13.4%) |
|  | Priority 3 (non-acute injury) | 1097 (7.8%) |
|  | Priority 4 (hold until verified need by another responding agency such as police or fire) | 30 (0.2%) |
|  | Priority Not Documented | 9040 (64.4%) |
| Urgency From Scene |  |  |
|  | Priority 1 | 805 (5.7%) |
|  | Priority 2 | 3770 (26.8%) |
|  | Priority 3 | 3589 (25.8%) |
|  | Priority Not Documented | 5863 (41.7%) |
| Skillset |  |  |
|  | ALS | 1535 (10.9%) |
|  | BLS | 12447 (88.7%) |
|  | Unknown | 5 (0%) |
| Languages |  |  |
|  | English | 11306 (84.7%) |
|  | Spanish | 782 (5.8%) |
|  | Russian | 377 (2.8%) |
|  | Cape Verdean | 337 (2.5%) |
|  | Haitian Creole | 72 (0.5%) |
|  | Cantonese | 62 (0.4%) |
|  | Portuguese | 32 (0.2%) |
|  | Mandarin | 32 (0.2%) |
|  | American Sign Language | 20 (0.1%) |
|  | Polish | 14 (0.1%) |
|  | Greek | 14 (0.1%) |
|  | Vietnamese | 13 (0.1%) |
|  | Italian | 12 (0.1%) |
|  | Hebrew | 11 (0.1%) |
|  | Persian | 10 (0.1%) |
|  | Other | 73 (0.5%) |

### Main Results

The matching algorithm, when evaluated on the training set (80%), had a sensitivity of 99.4%, positive predictive value of 99.9%, with an AUC of 0.99. When applied to the test set, it had an identical sensitivity of 99.4%, positive predictive value of 99.9%, with an AUC of 0.99. Model features and their weights are shown in [Table 3].

**Table 03:** Feature Weights

| Feature | n (%) | Coefficient | Odds Ratio |
| --- | --- | --- | --- |
| *dob_match* | 12750 (95.1%) | 2.8 | 16.9 |
| *lastname_match* | 12267 (91.5%) | 2.4 | 10.6 |
| *firstname_match* | 10728 (80%) | 1.9 | 6.9 |
| *first_name_jw* | n/a | 1.8 | 6.3 |
| *dob_levenshtien* | 12750 (95.1%) | 1.8 | 6.3 |
| *last_name_jw* | n/a | 1.8 | 6.2 |
| *ssn_match* | 8208 (61.2%) | 1.3 | 3.8 |
| *gender_match* | 13240 (98.8%) | 1.2 | 3.4 |
| *missing_ems_age* | 106 (0.8%) | 1.2 | 3.2 |
| *missing_ems_dob* | 104 (0.8%) | 1.2 | 3.2 |
| *missing_ems_firstname* | 49 (0.3%) | 1.0 | 2.6 |
| *missing_ems_ssn* | 4210 (31.4%) | 0.6 | 1.9 |
| *SSN_levenshtein* | n/a | 0.6 | 1.8 |
| *missing_ems_lastname* | 2 (0%) | 0.0 | 1.0 |
| *age_diff* | n/a | 0.0 | 1.0 |
| *missing_emr_age* | 0 (0%) | 0.0 | 1.0 |
| *missing_emr_dob* | 0 (0%) | 0.0 | 1.0 |
| *missing_emr_firstname* | 0 (0%) | 0.0 | 1.0 |
| *missing_emr_lastname* | 0 (0%) | 0.0 | 1.0 |
| *missing_emr_gender* | 0 (0%) | 0.0 | 1.0 |
| *missing_emr_ssn* | 0 (0%) | 0.0 | 1.0 |
| *missing_ems_gender* | 0 (0%) | 0.0 | 1.0 |

The feature *“dob_match”* had the highest odds ratio of 16.9 of predicting if an ePCR matched an EMR record followed by “lastname_match” (OR 10.6), *“firstname_match”* (OR 6.9) and *“firstname_jw”* (OR 6.3).

The inter-rater reliability between the primary and secondary reviewer was extremely reliable with a Kappa of 0.9.

## 4. Limitations

A significant limitation to our study is the reliance on a single EMS agency. Prior studies that have looked at multiple EMS agencies have demonstrated significant variations in practice among different companies.^3^ The use of a single agency could lead to a less generalizable algorithm due to less variability in data entry practices and quality.

Another limitation is the retrospective nature of our study which will need to be prospectively validated. While we were able to internally validate our probabilistic matching algorithm, its generalizability will be unknown until it is externally validated with different ePCR systems at different study settings.

## 5. Discussion

Prior studies have attempted to match prehospital records to inpatient records with varying degrees of success, reporting match rates ranging from 14% to 87%.^5,8^ Manual matching provides a higher success rate^8^ but is resource intensive, inaccurate,^9^ and slow. Most recently in 2015, Mumma successfully linked 34% of out of hospital cardiac arrest patients using an unsupervised probabilistic algorithm.^3^

To our knowledge, we are the first to propose a matching algorithm based on a supervised machine learning approach. In order to do so, we needed to collect gold standard labels, which was time consuming, but needed only to be done once. In Mumma’s work, they developed probabilistic algorithms using unsupervised techniques, using visual inspection of population statistics to help classify patients. Their unsupervised technique is limited by its assumption that all features are of equal importance, and does not allow for standard interrogation techniques of supervised models to uncover collinearity or non-linearity.

Having the ability to link ePCR with hospital records is crucial for improving patient care. Prior studies have demonstrated a lack of EMS documentation leading to increased mortality in trauma patients.^4^ A successful linkage algorithm would increase the amount of data available to hospital provides and potentially reduce trauma mortality. Another advantage to using a linkage algorithm is the increase in speed which the ePCR are available to be used by providers in the emergency department.

Linking the ePCR and hospital record allows hospitals to leverage prehospital care for both pay for performance metrics as well as publically available hospital performance reports. CMS rules allow hospitals to record interventions provided in the field, such as pain medication for long bone fracture or EKG acquisition and aspirin administration in non-traumatic chest pain, as having occurred during the hospital visit and occurring at time zero for process metrics provided the prehospital documentation is part of the hospital record. This has the potential to significantly improve reported metrics, and will have direct implications for reimbursement as payors continue the transition to quality payment programs. In addition to the direct implications on reimbursement, studies have shown publically reported measures have significant impact on hospital reputation and market share which further increases the potential financial benefits for hospitals and providers.^12,13^

When validating the model using the test set, there were 14 false negatives (0%) which were not matched by the algorithm. Three ePCR records were missing the first name, last name and DOB fields. The 11 remaining patients had inaccuracies with multiple fields such as name misspellings, incorrect first and last name, or incorrect DOB. These cases were matched by the reviewer based on a comparison of the EMS historical narrative and ED nursing triage note. Given the omitted/erroneous fields it is our impression that computer algorithm would not be able to match these patients without including additional data elements.

One result of this study is the ability to accurately and reliably merge EMS records with hospital records in real-time to create an integrated database. This allows us to monitor and research previously difficult prehospital questions such as on scene time in acute coronary syndrome^14^, or the impact of EMS interventions in trauma, or airway management in cardiac arrest.^15,16^

Failed or incorrect matches have the potential for significant impact on clinical care provided to the patient.^17^ Having a wrong patient matched could result in a patient receiving medications that they are allergic to or withholding medications because of a perceived allergy or prior administration. Providers also could believe that patient received treatment that was never administered. Our algorithm had both a high positive predictive value and negative predictive value that would result in very few failed or incorrect matches.

Future work will include natural language processing and semantic similarity analysis to compare the EMS historical narrative to the ED nursing triage note. We will also externally validate our algorithm at other sites with different EMS agencies utilizing different ePCR and hospital EMR platforms.

